# Insulin secretion chip (InS-chip) reveals two peaks within first-phase that temporally coincide with a transition in glucose metabolism

**DOI:** 10.1101/2023.03.07.531615

**Authors:** Yufeng Wang, Romario Regeenes, Mahnoor Memon, Jonathan V. Rocheleau

**Affiliations:** Advanced Diagnostics, Toronto General Hospital Research Institute, Toronto, Canada; Institute of Biomedical Engineering, University of Toronto, Toronto, Canada; Departments of Medicine and Physiology, University of Toronto, Canada

**Keywords:** Biological Sciences, Applied Biological Sciences

## Abstract

First-phase glucose-stimulated insulin secretion is mechanistically linked to type 2 diabetes yet the underlying metabolism driving this early stage of secretion is difficult to discern due to significant islet-to-islet variability. Here, we miniaturize a fluorescence anisotropy immunoassay onto a microfluidic device to measure C-peptide secretion from individual islets as a surrogate for insulin (InS-chip). This method measures secretion from up to four islets at a time with ∼7 s resolution while providing an optical window for real-time live cell imaging. Using the InS-chip, we reveal for the first time two glucose-dependent peaks of insulin secretion (i.e., a double peak) within the first phase (<10 min). By combining real-time secretion and live cell imaging, we show that islets transition from glycolytic to OxPhos-driven metabolism at the nadir of the peaks. Overall, these data validate the InS-chip to measure glucose-stimulated insulin secretion while revealing the first-phase secretion contains two peaks defined by a shift in glucose metabolism.

**Significance Statement:** Loss of the first phase of glucose-stimulated insulin secretion is one of the earliest signs of type 2 diabetes (T2D). Yet current strategies to measure the early dynamics are inadequate due to the need to pool secretion from multiple islets. In this study, we designed an islet-on-a-chip microfluidic device (InS-chip) to measure insulin secretion from individual islets with <7 s temporal resolution. Our design leaves an optical window for coupled live cell imaging to tease apart the metabolism underlying secretion. Our data reveal that first-phase insulin secretion is composed of two peaks that coincide with a shift in glucose metabolism.

## Text Introduction

Glucose stimulates a rise in islet b-cell ATP/ADP ratio resulting in a cascade of KATP channel closure, membrane depolarization, Ca^2+^-influx, and insulin secretion. However, this canonical model does not adequately describe why glucose-stimulated secretion (GSIS) occurs in a short 1^st^ phase burst within 5-10 min of stimulation followed by a smaller sustained 2^nd^ phase release [1], [2]. This biphasic pattern is found in humans under hyperglycemic clamp and perifused islets. The metabolism driving 1^st^-phase insulin secretion is particularly interesting since this phase is lost early in type 1 diabetes (T1D) [3] and type 2 diabetes (T2D) [4], yet difficult to explore due to the transient duration and islet-to-islet variability.

Humans and rodents show significant islet-to-islet variability in size and cellular composition [5], [6], oxygen consumption rate [7] and GSIS [8]. Hence, measuring GSIS from individual islets coupled with other live cell imaging techniques (e.g., Ca^2+^ and mitochondrial membrane potential (MMP) responses) is needed to explore the underlying mechanisms. Insulin secretion is commonly measured from hundreds of pooled islets using perifusion systems, which provides an accurate assessment of 1^st^- and 2^nd^-phase insulin secretion based on the timing and areas under the curve (AUC) [9], [10]. However, this method masks islet heterogeneity and generally cannot be used to simultaneously monitor other responses. More recently, various microfluidic devices have emerged to measure insulin secretion dynamically that also hold the potential for simultaneous islet imaging. However, many of these devices either have very low throughput (single islet loading) [11]–[13] or low sensitivity that requires pooled secretion from multiple islets [14]–[17]. Some of these devices sample islet effluent off-chip (e.g., Enzyme-linked immunosorbent assays (ELISA)) to achieve single islet sensitivity [13], [18], but this strategy suffers from uncoupling insulin secretion from other real-time responses. Off-chip analysis also requires additional washing steps, sampling system complexity, and costs that limit the application to specialized labs.

Fluorescence anisotropy immunoassays (FAIAs) are utilized extensively in clinical chemistry and bioassays to measure protein concentrations [19]. These assays are based on changes in fluorescence anisotropy due to the competitive binding of tracer (labelled peptide) and target to an antibody with the concentration simply quantified by the steady-state fluorescence anisotropy. Recently, FAIA was used on-chip to measure insulin secretion dynamically from both single islets [11] and pooled islets [14], [15]. Since insulin must dissociate from its hexamer crystal prior to the competition assay [20], these devices like many of the others use very long mixing channels (e.g., centimeter-long) to dissolve, mix, and reach the binding equilibrium. We postulate that dispersion due to these long channels ultimately diminishes the temporal resolution. Here, we adapted an FAIA to measure C-peptide from individual mouse islets in a hydrodynamic trap device (InS-chip). C-peptide is co-secreted with insulin in equimolar amounts but as a monomer, making it an ideal (i.e., fast mixing) surrogate for on-chip measurement of insulin secretion. We subsequently validated our method to measure GSIS from individual islets with ∼7 s resolution, which allowed us to detect two peaks of secretion inside the 1^st^ phase response. Finally, the simple design allowed us to combine real-time secretion and live cell imaging to show a transition from glycolytic-to mitochondrial metabolism associated with a transition from the first peak to the second peak of 1^st^-phase GSIS.

## Results

### Development and characterization of FAIA

The dynamic range of FAIAs depends on the difference in molecular size between the unbound and antibody-bound sensor. To lower the unbound anisotropy and effectively maximize the dynamic range of our FAIA, we designed an N-terminally tagged C-peptide sensor (C-peptide*) based on 10-amino acid residues that encompassed the antibody epitope that was only ∼1/3^rd^ the size of full-length C-peptide. The sensitivity of FAIAs is primarily determined by equilibrium constants for direct binding between the C-peptide* and Ab (K_D1_), and competition between the pre-bound C-peptide*-Ab complex and endogenous C-peptide (K_D2_) [21] (**Fig. 1A**). To measure K_D1_, Ab was serially diluted in 100 nM C-peptide* and the anisotropy response was fit to the direct binding curve (eq. 3-4) to yield a value of 180 ± 12 nM (**Fig. 1B**). A C-terminally tagged C-peptide* showed higher affinity (K_D1_ = 2.89 ± 1.44 nM) (**Fig. S1**), but we chose the N-terminally tagged C-peptide* since a higher K_D1_ is often the result of a higher dissociation rate (k_off_) [22], which would shorten the time to reach equilibrium on-chip [23]. To measure K_D2_, C-peptide was serially diluted in pre-mixed Ab-C-peptide* and the anisotropy response was fit to the competitive binding curve (eq. 3, 5-8) to yield a value of 5.8 ± 2 nM (**Fig. 1C**). Notably, the relative standard deviation (RSD) of the data was 4.4 ± 0.62 %, suggesting that the assay could be applied to track relative changes, but would be less reliable in the quantification of absolute C-peptide levels. In addition, a parameter sweep of varying secretion concentrations was performed to determine the concentrations of C-peptide* and Ab for maximum assay sensitivity (**Fig. S2**). Independent of C-peptide concentration, maximum sensitivity is achieved when Ab concentration is kept at the K_D1_ value, consistent with previous work [24]. We chose to set the C-peptide* concentration to 100 nM for all experiments since it gives a 10-fold higher signal than the background (data not shown), ensuring assay sensitivity without impacting signal variability [25], [26].

**Fig. 1.**
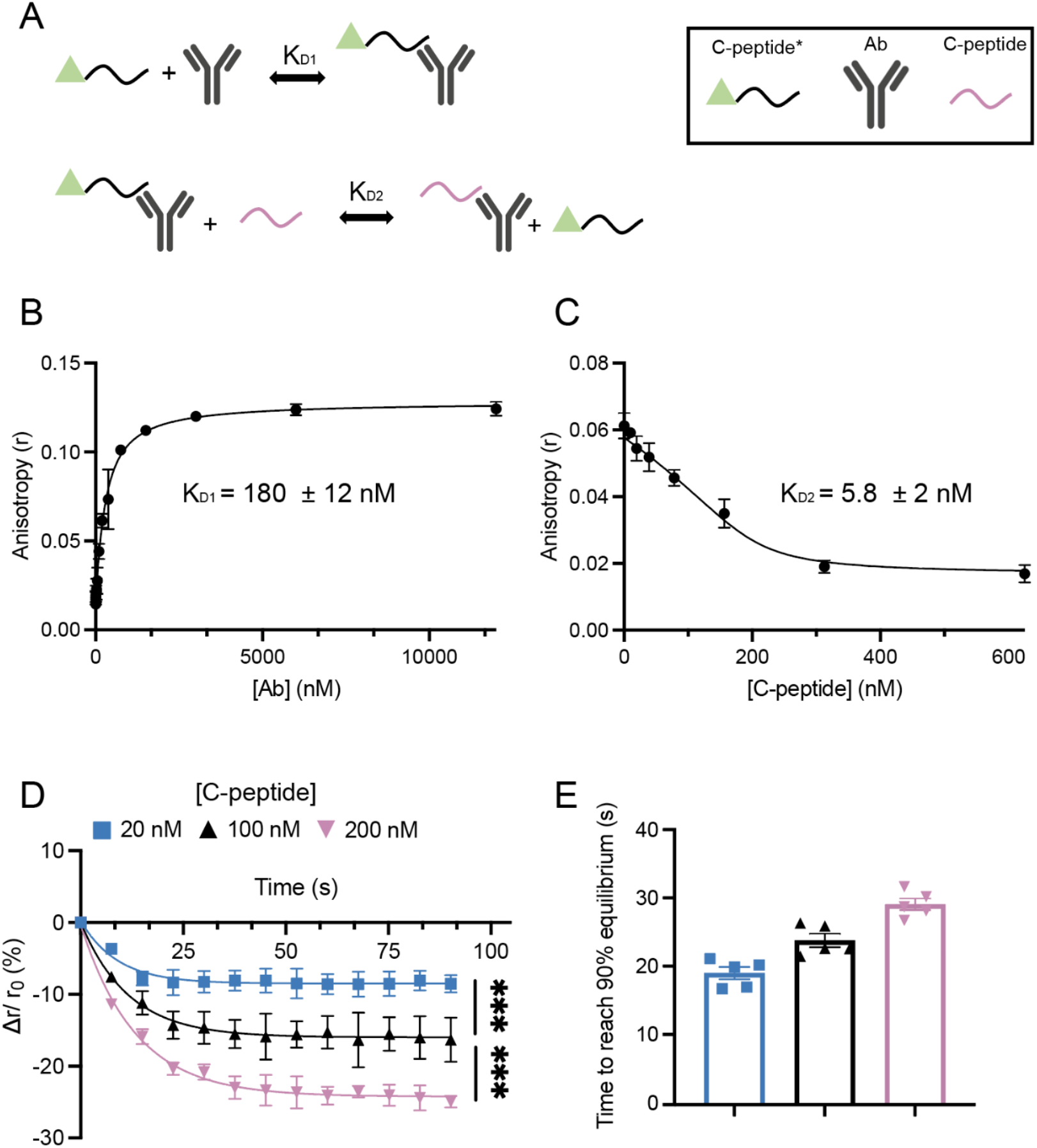
Development of an FAIA for the detection of insulin secretion using a C-peptide-based sensor (C-peptide*). (**A**) Schematic showing direct binding (top) and competitive binding (bottom) for the characterization of K_D1_ and K_D2_, respectively. (**B**) Fitting of the direct binding curves between serially diluted Ab and 100 nM C-peptide* yielded a K_D1_ of 180 ± 12 nM (n = 3). (**C**) Fitting of competitive binding curves between serially diluted C-peptide and pre-mixed 200 nM Ab and 100 nM C-peptide* yielded a K_D2_ of 5.8 ± 2 nM (n = 3). (**D**) Kinetics of competitive binding between different concentrations of C-peptide and pre-mixed 200 nM Ab and 100 nM C-peptide* in the device shown in **Fig S3A** (n = 5 for each c-peptide concentration). Anisotropy values after 30 s of competition with the indicated concentration of C-peptide. *** indicates p < 0.001 by one-way ANOVA. (**E**) Time taken to reach 90% plateau of the competition kinetics curves shown in (**D**).

Our goal was to measure secretion on-chip yet FAIAs require time to reach equilibrium. To determine the on-chip residence time required to reach equilibrium, we built a continuous-flow microfluidic that quickly mixes the solutions using sheath-flow mixing in a straight channel, followed by a serpentine channel to provide extended residence time for the assay to equilibrate (**Fig. S3**) [27]. We filled the middle inlet of the device with varying C-peptide (20 nM, 100 nM, 200 nM) and the outer two inlets with pre-mixed 100 nM C-peptide* and 200 nM Ab. The device achieved mixing in less than 1 s and provided a long residence time (> 85 s) for the assay to equilibrate (**Fig. S3B**). The progress of competitive binding C-peptide (20, 100, and 200 nM) was imaged at different locations along the microfluidic device (**Fig. 1D**). These data show that as unlabeled C-peptide displaces C-peptide* from the Ab the anisotropy lowers, reaching 90% of the equilibrium within 19- to 29-s depending on the amount of C-peptide (**Fig. 1E**). In contrast, the high-affinity C-peptide* required nearly twice the time with 200 nM C-peptide to reach equilibrium (**Fig S1C-D**). Overall, these data show that assay sensitivity is optimized by setting Ab concentration to the K_D1_ value (200 nM) and that the assay requires ∼29 s on-chip to reach equilibrium.

### Design of islet-on-a-chip

We previously designed several islet-on-a-chip devices to treat and image living pancreatic islets [28]–[30]. To measure insulin secretion from individual islets, we aimed to adapt one of our islet-on-chip devices for the FAIA of C-peptide (**Fig. 2**). Microfluidic devices are generally fabricated using PDMS due to biocompatibility, oxygen-permeability and standard fabrication that translates to other research laboratories [31]. We generally bond our islet-on-a-chip devices to No. 1.5 glass coverslips to hold the islets directly against a glass thickness optimized for high-resolution fluorescence imaging. The device shown is filled with food coloring to illustrate the islet-inlet/outlet tubes and the 4 microfluidic islet chambers (**Fig. 2A**). By placing this device on a fluorescence anisotropy microscope, our goal was to flow Ab-bound-C-peptide* past islets trapped in the device using a single vacuum-based pressure controller (Fluigent, MA) and image C-peptide secretion based on anisotropy of the competition assay.

**Fig. 2.**
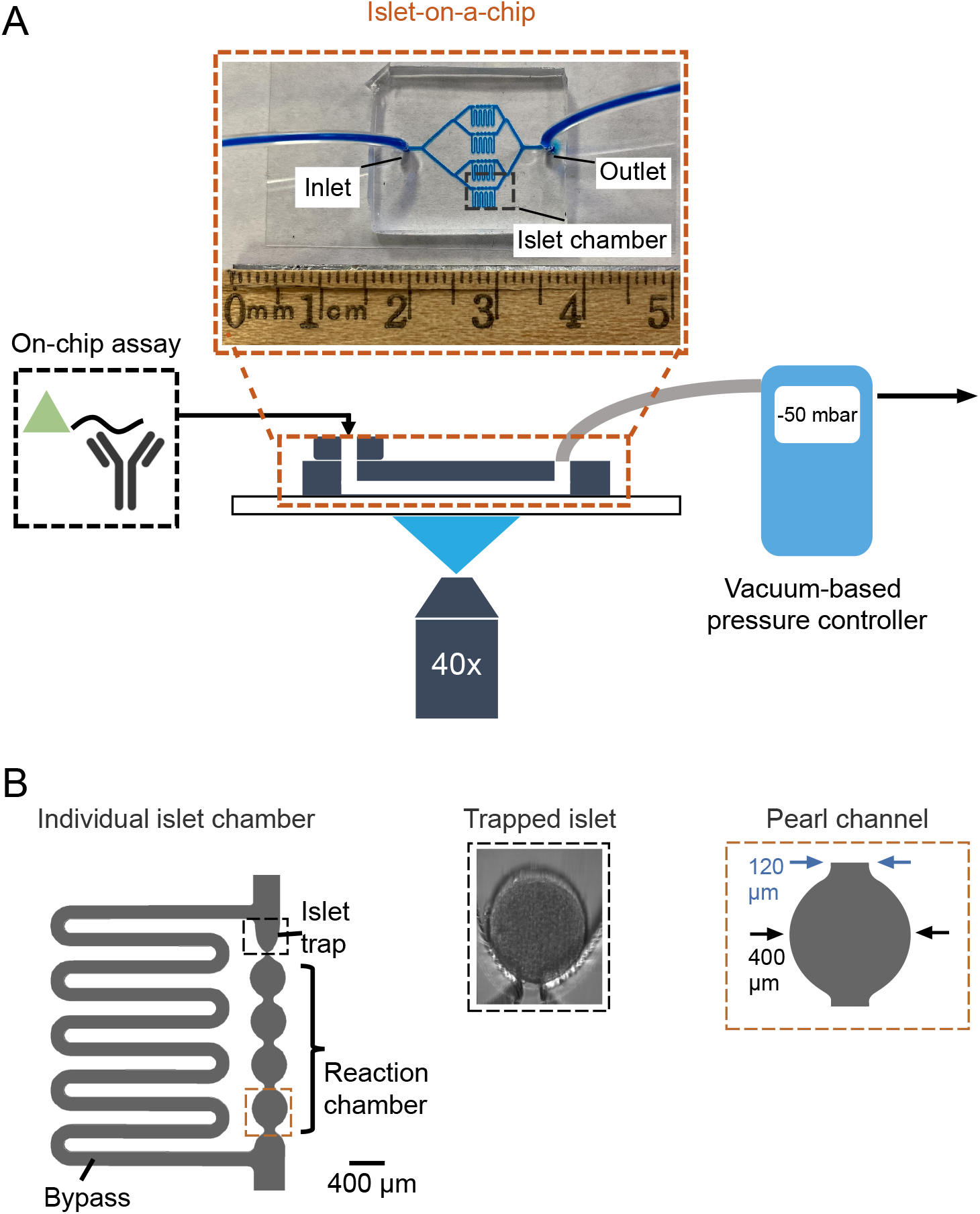
Design of an islet-on-a-chip to hold four individual islets for imaging insulin secretion. (**A**) Schematic of the microfluidic device mounted on a microscope stage and reagents drawn through the device using a vacuum-based pressure controller. A photograph of the device filled with blue dye is shown for easy visualization of the microfluidic channels. In the device shown, the inlet tubing was used instead of the on-chip reservoir to prevent obstruction of the channel features. Flow in the device was controlled by a single vacuum-based pressure controller. (**B**) Schematic showing one islet chamber (left) consisting of a hydrodynamic trap to stabilize the tissue during imaging (middle). This trap chamber includes a bypass channel to prevent pressure buildup against loaded islets and a reaction chamber consisting of a channel with sequential pearl shapes varying from 120-to 400-mm in width (right).

We incorporated four hydrodynamic traps to hold the islets separately in the device [28](**Fig. 2B**). These traps were modified to increase the residence time of effluent on the chip to allow the assay time to reach equilibrium. (**Fig. 2B, left**). This included elongating the bypass and pearl-shaped reaction channels. The lengths of the bypass and reaction channels were matched to ensure easy loading of islets into the hydrodynamic trap (R_pearl_ < R_bypass_) (**Fig S4A-B**). To confirm that a longer bypass channel was not increasing shear stress, we modelled an islet under a flow rate of 50 μL·h^-1^ (**Fig. S4C-D**). These data show that most of the shear stress on the islet surface is less than 10 mPa (**Fig S4D**) like our previous design [28]. In hydrodynamic traps, the islet effluent flows through the nozzle and down the reaction channel (**Fig. 2B, middle**). Once an islet fills a trap, most of the flow into the chamber goes down the bypass channel, which effectively prevents a pressure buildup against the islet (i.e., minimizes mechanical stress) and slows flow past the islet (i.e., reduces shear stress) [32]. The slow flow past the islet ensures tissue function while minimally diluting the effluent. The effluent of each islet enters an independent reaction chamber consisting of a series of pearls that vary in width periodically from 120-to 400-μm (**Fig. 2B, left and right**). As will be shown, the pearl-shaped reaction chamber allows islet secretion to mix and reach equilibrium with the C-peptide*-Ab complex and provides fiducial marks for imaging. The effluent from each islet subsequently merges with the bypass channel into a single outlet for flow in the four chambers to be controlled by a single vacuum-based pressure controller. Overall, this new design holds 4 islets in well-defined parallel hydrodynamic traps that collect the effluent in independent reaction chambers.

### Evaluation of mixing efficiency

Media flows both around and through islets in hydrodynamic traps [28]. Thus, we were concerned that media pushed through the islet (i.e., Ab-free effluent) would need time to mix with the carrier media prior to the competitive binding. To evaluate the speed of mixing in the pearl channel, we designed a microfluidic device to mimic effluent mixing with carrier media by merging three inlets into a single pearled channel (**Fig. 3A**). We placed FITC-dextran in the outer inlets to mimic Ab in the carrier media (∼150 kDa) and imaged the mixing efficiency of clear effluent from the middle inlet along the channel (**Fig. 3B**). These data show greater mixing efficiency at all positions along the channel and with lower flow rates, consistent with greater residence time increasing lateral diffusion-based mixing (**Fig. 3C**). At 25 μL·h^-1^, the mixing efficiency near the device outlet (i.e., the 9^th^ pearl) exceeded 90% (**Fig. 3C**). To evaluate the mixing time relative to the equilibration time of the FAIA, we next flowed 200 nM C-peptide in the outer stream and 100 nM Ab-C-peptide* in the inner stream and followed the mixing efficiency relative to the change in anisotropy with time down the channel (**Fig. 3D-E**). The mixing efficiency plateaued before 10 s while the anisotropy was still dropping at 20 s (**Fig. 3D**). Based on fits to these curves, the time required to reach the 90% plateau for mixing and competitive binding was ∼3 and 29 s, respectively (**Fig. 3E**). These data show the kinetics of mixing is much faster than the on-chip FAIA. More importantly, the slowest flow rate we could reach empirically due to pump instability (25 μL·h^-1^) was 5′ faster than the flow rate around islets measured by PSV (∼ 5 μL·h^-1^, **Fig. S5**). To evaluate mixing efficiency at a 5 μL·h^-1^, we modelled the diffusion of a species in the outer two inlets of the same device with a diffusion coefficient approximating an antibody (2.83 × 10^−11^ m^2^s^-1^) [14] (**Fig. 3F**). This modelling suggests the mixing efficiency reaches >90% within 1 s. In addition, at 5 μL·h^-1^, the first pearl channel provides a top-to-bottom residence time of 26 s, which is close to the time required to reach 90% equilibrium for FAIA on-chip (**Fig. 1E**). Nonetheless, our final channel design includes 4 pearl channels to ensure adequate residence time to mix and reach equilibrium. Overall, these data demonstrate that mixing is much faster than the kinetics of competitive binding in our device.

**Fig. 3.**
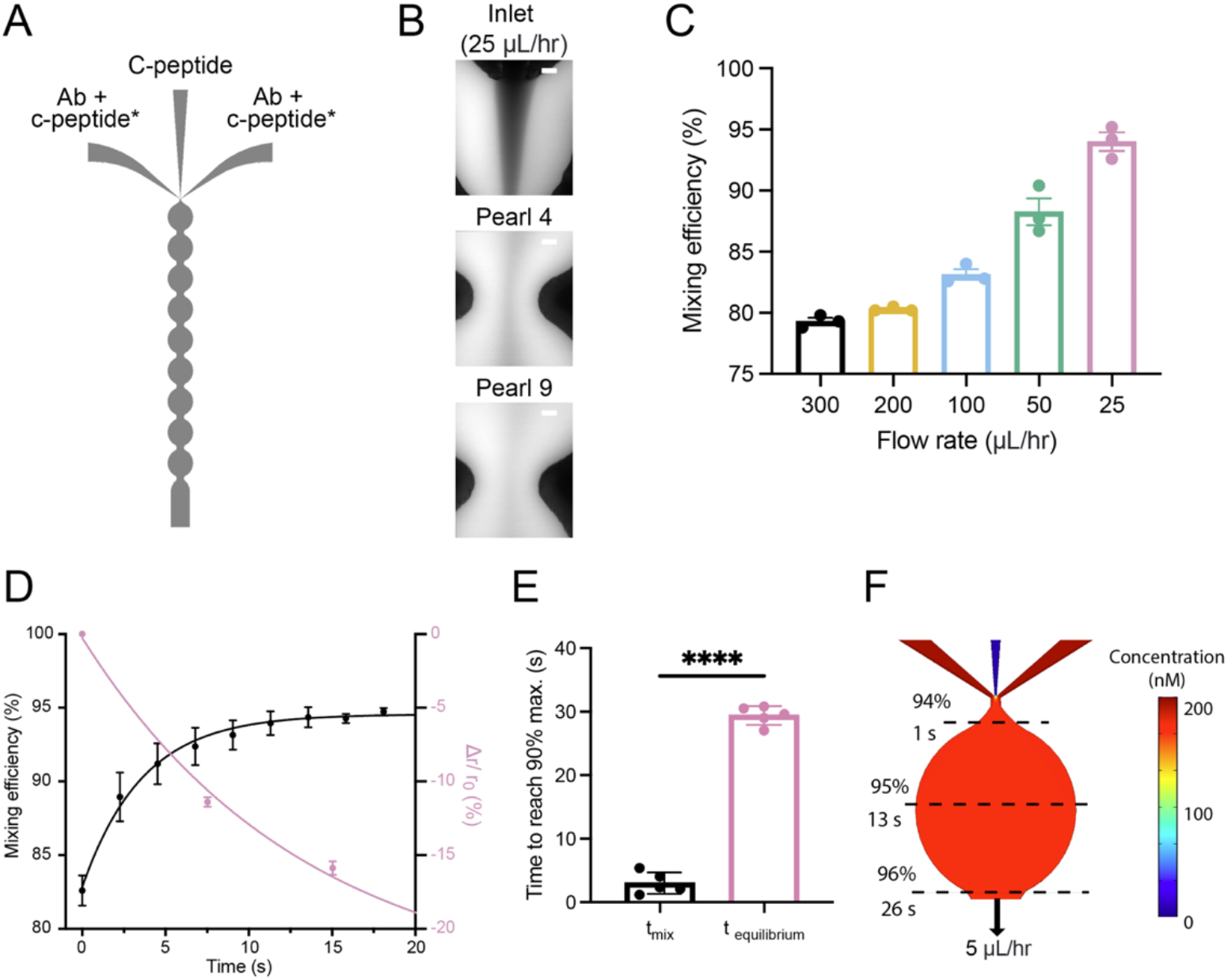
Characterization of on-chip mixing reveals assay equilibrium is rate-limiting. (**A**) Schematic of the microfluidic device used to evaluate mixing on-chip. The three-channel inlet simulates media flowing around-and through islets by flowing media containing pre-mixed Ab and c-peptide* in the outer channels to sandwich secreted C-peptide in the middle channel. (**B**) Fluorescent images of the inlet, 4^th^ and 9^th^ pearl channel when 200 nM FITC-Dextran is drawn from outer channels and imaging buffer is drawn from the middle channel. Scale bars represent 25 μm. (**C**) Mixing efficiencies were obtained at the 9^th^ pearl channel at varying flow rates from the outlet (n = 3 for each flow rate). (**D**) Changes in the mixing efficiency (left y-axis) and change in anisotropy due to competition with 200 nM C-peptide in the first 20 s (right y-axis). (**E**) Comparison of the time to reach the 90% plateau of the mixing curve (t_mix_) and competition curves (t_equilibrium_) (**D**). **** indicates p < 0.0001 by unpaired t-test. (**F**) COMSOL simulation of the first pearl of the on-chip mixing device shown in (**A**) at 5 μL/hr. The diffusion coefficient of the species in outer inlets was set to 2.83 × 10^−11^ m^2^/s to simulate an Ab. The dashed lines indicate residence times of 1-, 13- and 26-s, corresponding to 94%, 95% and 96% mixing efficiencies, respectively.

### Characterization of residence time and temporal resolution

Our goal was to measure the temporal dynamics of insulin secretion that likely occur in pulses/bursts due to metabolically driven Ca^2+^ activity [33]. Long microfluidic channels suffer from dispersion such that a pulse of solute would disperse flowing down a channel [34]. Thus, we were concerned that dispersion along the pearl-shaped reaction channel would dampen these dynamics. To evaluate the impact of residence time (t_residence_) on the dispersion of a pulse, we rapidly switched the solution in the on-chip reservoir from a blank buffer to FITC-dextran to create a step in the dye concentration and imaged the change in intensity temporally at different points along the chamber (**Fig. 4**). We anticipated this stepped change in concentration would disperse through the entire device, but we were particularly interested in measuring the dispersion from the islet through the pearl channel to determine how a pulse of insulin would be dispersed (**Fig 4A**). The fluorescence intensity increased at the islet trap before the pearls and slowly ramped with time in both regions consistent with dispersion throughout (**Fig 4B-D**). Plotting the intensities at the trap relative to the 1^st^ (**Fig. 4C**) and 4^th^ (**Fig. 4D**) pearls showed significant dispersion of the dye at each position along the pearl with a progressively shallower slope. This difference in dispersion becomes more apparent once the signal between the trap and pearls are temporally aligned and the difference in intensity is plotted (**Fig 4E**). To quantify the temporal resolution at different pearls of the channel, we calculated the differences in time to reach 90% of saturation (t_90%-10%_) between the islet trap and either the 1^st^ or 4^th^ pearl (**Fig 4F-G**) [15]. These data show the 1^st^ pearl provides a residence time of 29 s with a temporal resolution of 6.8 s. In contrast, the 4^th^ pearl provided a residence time of 85 s but with a significantly lower temporal resolution of 54 s. Overall, these data suggest that imaging at the 1^st^ pearl provides superior temporal resolution while still providing sufficient residence time to reach the equilibrium time of the FAIA.

**Fig. 4.**
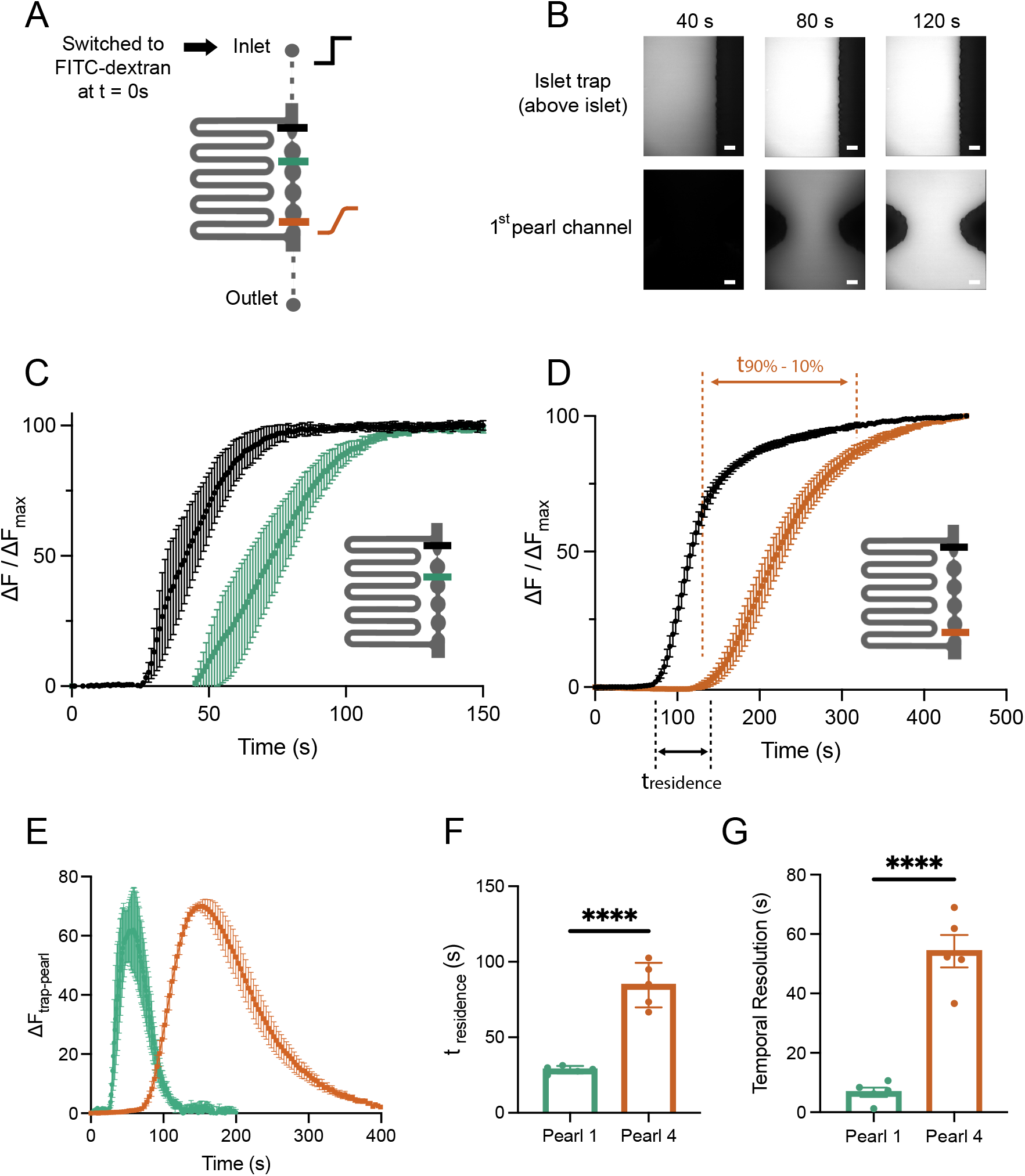
Evaluation of dispersion in the islet-on-a-chip device. (**A**) Schematic of the experimental setup for characterization of dispersion. At time 0, the solution at the inlet was switched from imaging media to media containing 200 nM FITC-Dextran (step increase in black). As it travels through the device, the signal becomes more dispersed (sigmoidal increase in brown). Fluorescence was imaged simultaneously at the trap region (black line) and 1^st^ (green line) and 4^th^ pearls of the channel (brown line). (**B**) Representative fluorescent images of the islet trap and 1^st^ pearl channel taken 40-, 80-, and 120-s after the step increase at the inlet. Scale bars represent 25 μm. (**C-D**) Normalized changes in fluorescence intensity in the islet trap, and 1^st^ (**C**) and 4^th^ pearl (**D**) (n = 5 for each pearl). The residence time (t_residence_) is defined as the time taken for fluorescence to reach 10% of the maximum between the two traces in each graph. The 90% recovery time (t_90%-10%_) is defined as the time taken for fluorescence to rise from 10% to 90% of the maximum intensity. (**E**) The average fluorescence difference spread from the islet trap to the 1^st^ and 4^th^ pearls. (**F**) Comparison of t_residence_ between the 1^st^ and 4^th^ pearl (n = 5). **** indicated p < 0.0001 by unpaired t-test. (**G**) Comparison of temporal resolution achieved by imaging at 1^st^ and 4^th^ pearl (n = 5). **** indicated p < 0.0001 by unpaired t-test.

### Imaging GSIS in individual pancreatic islets

To validate the ability of our InS-chip to measure GSIS dynamically in individual islets, we measured the glucose-stimulated response of single islets (4 at a time) isolated from FVB mice by measuring the FAIA response in the 1^st^ pearl (**Fig. 5A-G**). As mentioned previously, the FAIA assay showed a variance appropriate for measuring relative changes in the competitor rather than absolute values. Hence, here we report the relative anisotropy response to reflect the dynamic changes in insulin secretion. Shown are representative traces of individual islets responding to 8-(**Fig 5A**), 11- (**Fig 5B**), and 20-mM glucose (**Fig 5C**) followed by treatment with 100 mM tolbutamide (**Fig. 5A-C**) or diazoxide (**Fig. 5D**). All these curves show relative anisotropy responses characteristic of a 1^st^ phase rise (<10 min), followed by a lower 2^nd^ phase. Interestingly, 10 of the 44 traces showed visible oscillations in the 2^nd^ phase (e.g., **Fig 5D**) with periodicities between 0.8 and 3 min. Consistent with GSIS, the responses were increased by the K_ATP_ channel blocker (tolbutamide) (**Fig 5A-C**) and decreased by the activator (diazoxide) (**Fig 5D**). Comparing these responses across several islets showed a significant tolbutamide-induced increase and diazoxide-induced decrease, respectively (**Fig 5E**). Similar analysis comparing the 8-mM glucose-stimulated responses to 11- and 20-mM glucose responses showed a significant increase; however, we did not observe a significant difference between the 11- and 20-mM glucose-stimulated responses. Finally, analysis of individual islets stimulated with 11 mM glucose showed significant heterogeneity with no correlation between islet size and either 1^st^ or 2^nd^ phase GSIS (**Fig 5F-G**). This may reflect a limitation in the design of the holding chamber as it was not designed to accommodate all sizes of islets. Nonetheless, these data reiterate the power of the device lies in measuring the timing and shape of secretion rather than in comparing absolute islet-to-islet secretion variability. To validate the ability of the device to couple GSIS with live cell imaging, we loaded islets with Cal-520 and simultaneously imaged cytoplasmic Ca^2+^ and insulin secretion (**Fig 5H-I**). The Ca^2+^ images were collected from the islets in each trap while the insulin response was imaged at the 1^st^ pearl (**Fig 5H**). This setup allowed us to compare the temporal responses to 11 mM glucose followed by subsequent membrane depolarization by 30 mM KCl (**Fig. 5H-I**). The representative Ca^2+^ trace shows the characteristic spikes in secretion associated with simultaneous Ca^2+^-influx through voltage-gated channels induced by glucose and subsequent membrane depolarization by KCl (**Fig 5I, green trace**).

**Fig. 5.**
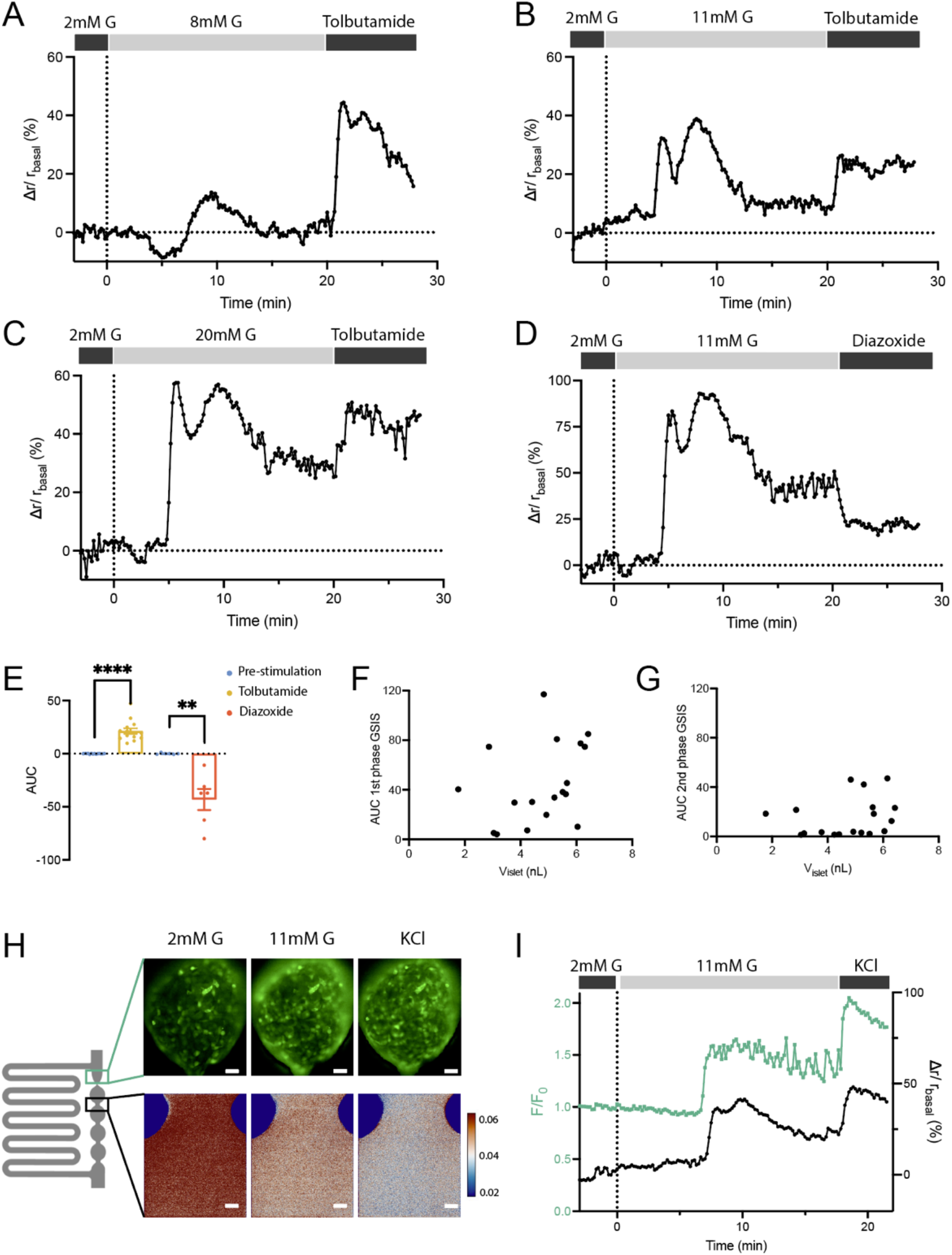
Validation of the Insulin Secretion-chip (InS-chip) to measure GSIS of individual murine islets. (**A-C**) Representative traces of insulin secretion as individual islets were treated with 2 mM glucose followed by 8- (**A**, n = 13), 20- (**B**, n = 15) and 11-mM glucose (**C**, n = 16), and 100 μM tolbutamide as indicated. The change in anisotropy of C-peptide* (Dr) was normalized to the anisotropy at 2 mM glucose (r_basal_). (**D**) Representative traces of insulin secretion as individual islets were treated with 2- and 11-mM glucose followed by 100 μM diazoxide (n = 6). (**E**) The average area under the curve (AUC) of GSIS traces when islets were treated with tolbutamide (**C**) or diazoxide (**D**) at 11 mM glucose. (**E**) Individual traces were normalized to 15 data points prior to the addition of tolbutamide and diazoxide. ** indicated p < 0.01, **** indicated p < 0.0001 by paired t-test. (**F-G**) AUC of 1^st^ phase (**F**) and 2^nd^ phase (**G**) 11 mM glucose-stimulated responses relative to islet volume (n = 18). (**H**) Representative images of a mouse islet loaded with Cal-520 (top) treated with 2- and 11-mM glucose followed by 30 mM KCl. The 1^st^ pearl was simultaneously imaged for anisotropy to measure secretion (bottom). Scale bars represent 25 μm. (**I**) Representative trace of Ca^2+^ response and relative change in anisotropy (n = 8).

Notably, insulin secretion and Ca^2+^ activity were both largely in sync throughout (**Fig. 5I, black trace**). Overall, these data suggest our InS-chip can dynamically track insulin secretion from individual islets while providing an optical window to couple with other live cell imaging techniques. We used this window to confirm the synchrony of insulin and Ca^2+^ responses, further validating our ability to trace insulin secretion in individual islets.

### Single- and Double-Peak 1^st^ Phase GSIS

GSIS measured from pooled islets is conventionally defined by a 1^st^-phase spike in insulin secretion that occurs within 10 min of a step increase in glucose followed by a lower 2^nd^ phase [35]. Critically, a loss of 1^st^ phase insulin secretion is an early sign of T1D and T2D and is thus mechanistically linked to both diseases [3], [4]. To explore the dynamics of the 1^st^ phase response, we stimulated islets in our device with 8-, 11-, and 20-mM glucose and focused our analysis on the first 20 min of stimulation (**Fig 6A-C**). The two representative traces demonstrate the significant variability in the time-to-turn-on and the shape of the curves even when islets were stimulated with the same concentration of glucose (**Fig. 6**). Variability in the time-to-turn-on is consistent with similar variability in islet Ca^2+^ responses [36]. The traces also consistently revealed either a single-peak (**Fig. 6A, black trace and Fig. 5A**) or a double-peak (**Fig. 6A, marron and Fig. 5B-D**). A double peak occurred only when stimulating with ≥ 11 mM glucose (**Fig. 6A and Fig. 5B-D**) with a clear glucose-dependent prevalence with ∼58% of islets showing a double peak at 11 mM glucose and > 85% at 20 mM glucose (**Fig. 6B**). We subsequently compared the timing of the single and double peak maxima (**Fig. S6**). These data show that the single-peak responses reach a maximum at the same time as 2^nd^ peak when present. In addition, insulin secretion started earlier when a double peak was present (**Fig. 6C**). These data suggest that islets showing a double peak are metabolically more active, perhaps reflecting islet functional heterogeneity. To investigate the metabolism underlying a double-peak response, we took advantage of the optical window underneath the islets to simultaneously measure GSIS (**Fig. 6D-G**) while imaging MMP using TMRE (**Fig. 6D-E**) and ATP using PercevalHR (**Fig. 6F-G**). We stimulated with 20 mM glucose to induce double-peak secretion followed by 2 mM FCCP to uncouple MMP. In the representative traces, TMRE shows a relatively slow rise in response to glucose and a steep fall in response to FCCP, consistent with changes in MMP [37] (**Fig. 6E**). On the other hand, ATP rose in response to glucose more quickly reaching a plateau followed by a small dip and recovery, consistent with previous findings [38]. The corresponding insulin traces showed a double-peak response to glucose albeit somewhat less prominently in islets loaded with TMRE. We postulate that the subdued response may be due to the entry of this cationic dye into the mitochondrial matrix, impacting MMP and respiration [39]. To further explore the double-peak response, we combined the responses of multiple individual islets by aligning the MMP (**Fig. 6H**) and ATP curves (**Fig. 6I**) to three critical time points of insulin secretion highlighted in Fig. 6E and G. The first point (Point i) represents the switch from low to high glucose prior to the 1^st^ phase insulin response (**Fig. 6H and I, left**). These data show a small elevation in the MMP response within 3 min of glucose stimulation compared to a much larger rise in ATP. These data are consistent with glycolysis-driven ATP production [40], [41]. The second point (Point ii) represents the initiation of insulin secretion (**Fig. 6H and I, middle**). These data show that upon initiation of insulin secretion, MMP slowly ramps up, reaching slightly above 50% of the maximum response. Meanwhile, ATP, which had previously plateaued, starts to dip. These data are consistent with a transition from glycolysis-driven metabolism to activation of OxPhos. The final point (Point iii) represents the transition between the two insulin peaks (i.e., the nadir) (**Fig. 6H and I, right**). These data show that as the second peak develops the MMP rises continuously. In contrast, ATP starts to rise again only at the 2^nd^-peak of insulin secretion.

**Fig. 6.**
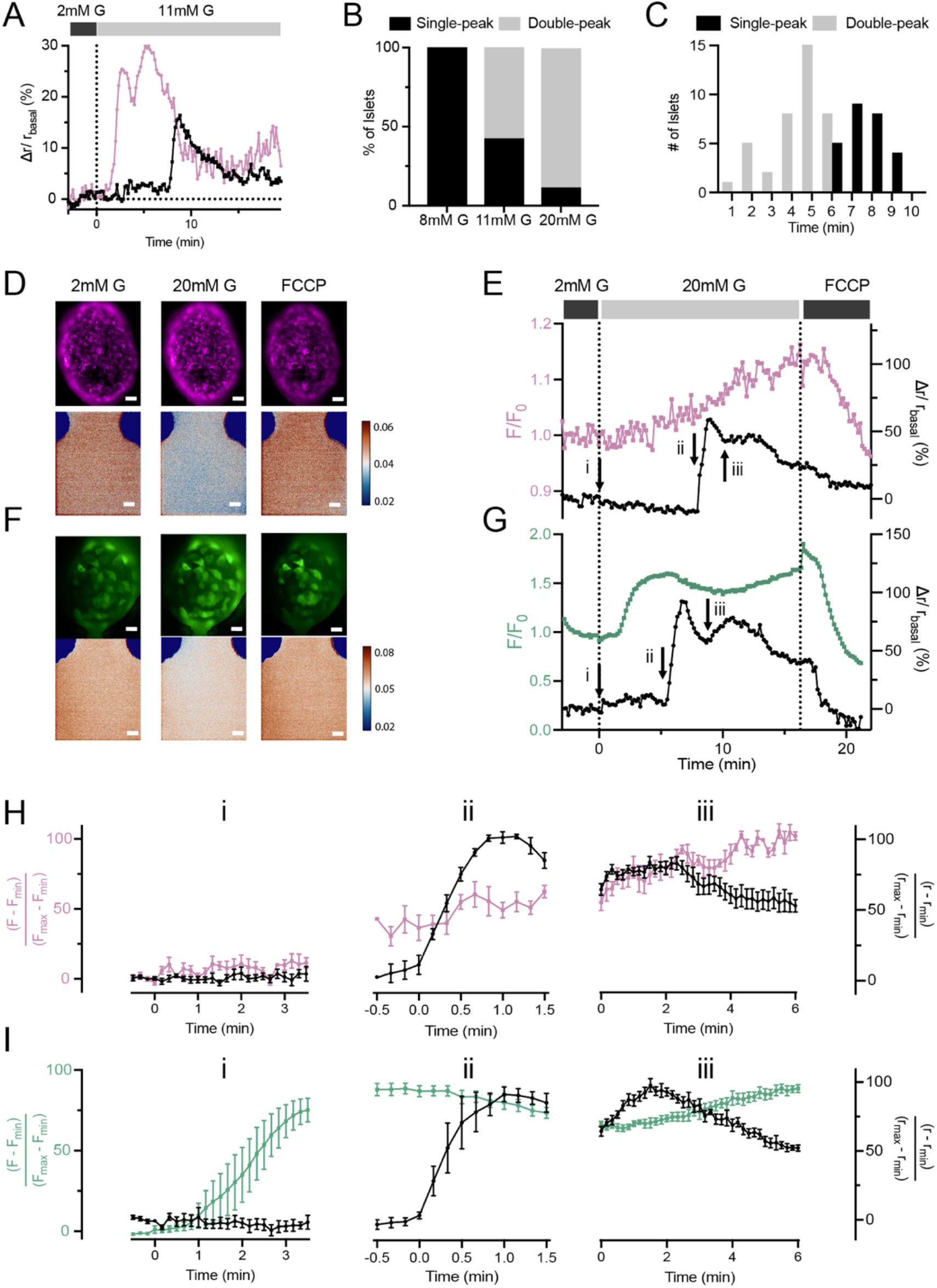
Identification and characterization of 1^st^ phase responses showing single- and double-peak GSIS. (**A**) Representative traces of two different islets stimulated with 11 mM glucose demonstrating a single- and double-peak in the 1^st^ phase response (<10 min of stimulation). (**B**) The percentage of islets with single- or double-peak GSIS after stimulation with 8- (n = 13), 11- (n = 16) and 20-mM glucose (n = 15). (**C**) Frequency distribution of the response times of 1^st^ phase GSIS after stimulation with 8-, 11- and 20-mM glucose. (**D, F**) Representative images of a mouse islet loaded with TMRE (**D**, *top*) or PercevalHR (**F**, *top*) and 1^st^ pearl for simultaneous imaging of insulin secretion (**D&F**, *bottom*). Scale bars represent 25 μm. (**E, G**) Representative traces of GSIS with TMRE (**E**) and PercevalHR (**G**) from individual mouse islets. Both the TMRE and PercevalHR responses were normalized to fluorescence intensity at 2 mM glucose. Black arrows indicate “critical time points” of insulin secretion for further analysis. (**H-I**) The average indexed changes in GSIS with TMRE (**H**) and PercevalHR (**I**) was determined by setting time = 0 min at the indicated time points shown in (E, n=5) and (G, n=5). Point i represents the switch from 2- to 20-mM glucose, point ii represents the start of 1^st^ phase insulin secretion, and point iii represents the low between the double-peaks of 1^st^ phase GSIS.

Consistent with other work, we postulate this dip and recovery of ATP represents greater consumption due to the “work” of insulin secretion followed by Ca^2+^-activated mitochondrial metabolism [38]. Overall, these data suggest 1^st^-phase GSIS is composed of a double-peak response involving a transition from glycolysis- to mitochondrial-dominant metabolism.

## Discussion

We developed a real-time assay for C-peptide secretion from individual islets with simultaneous live-cell imaging (i.e., InS-chip). Previous microfluidic devices designed to measure insulin secretion can measure responses from a small number of islets and even from individual islets, yet these devices are considerably more complex and/or lack the ability to correlate secretion with other metabolic readouts. The InS-chip measures secretion from 4 individual islets with ∼7 s resolution using a single vacuum pump and modified fluorescence microscope. We achieved this resolution through a number of design choices including targeting C-peptide rather than insulin, selecting a low-affinity antibody-tracer pairing, and imaging as close to islets as the assay would allow in the hydrodynamic traps. We validated the InS-chip could measure mouse islet secretion in response to glucose and modulators of K_ATP_ channel activity, and by simultaneous Ca^2+^ imaging. Our data reveal two glucose-dependent peaks of insulin secretion inside what is classically described as 1^st^-phase insulin secretion (<10 min). We subsequently used the InS-chip to correlate insulin secretion with changes in MMP and ATP. These data revealed a transition from glycolytic-to OxPhos-dependent insulin secretion at the nadir of the two peaks of insulin secretion. Overall, this method provided exceptional temporal resolution of secretion that allowed us to see new details of 1^st^-phase insulin secretion (single versus double-peaked secretion) that correlate with a transition in glucose-stimulated metabolism.

Many recent islet-on-a-chips pool secretion from multiple islets [14]–[17], which would mask islet heterogeneity and blur temporal dynamics. Our ability to track individual islets showed variability in the time-to-turn on and pattern of secretion (e.g., single versus double peak) even under the same level of glucose stimulation. Previous devices capable of measuring insulin secretion from individual islets either have had relatively low throughput (single-islet loading) [11]–[13] or required off-chip quantification, notably through ELISA [13], [18]. Capillary electrophoresis has been used to measure insulin secretion from multiple, individual islets on-chip [42]. This method provides high temporal resolution and sensitivity but requires a laborious device manufacturing process and a complex experimental setup (e.g., external power supply and relay systems), making it challenging for other laboratories to adapt. In contrast, our all-optical assay on-chip allows real-time insulin secretion using a single vacuum pump and a widefield microscope modified for fluorescence anisotropy imaging.

Our InS-chip achieved a high ∼7 s temporal resolution due to the careful design of the assay and microfluidic device. First, microfluidic mixing generally relies on diffusion due to laminar flow.

Accordingly, many islet-on-a-chip devices integrate long mixing channels (often cm in length) between the islet and detection area. In contrast, our results show that the time required for mixing was negligible compared to the FAIA equilibrium time. This was achieved first by incorporating the Ab-C-peptide* complex in the media such that the small volume of media that went around the islet immediately surrounded/enveloped the effluent. Second, we used hydrodynamic traps that send relatively slow flow past the islet (∼ 5 mL·h^-1^). This limits the dilution of islet effluent and allows for longer residence time in the pearled channel [28]. Third, we designed our FAIA to measure C-peptide as a surrogate for insulin. Endogenous insulin is secreted as a hexamer crystal with Zn^2+^ that would need to dissolve prior to the competition assay [20]. C-peptide is instead secreted as a monomer, but in an equimolar ratio to insulin. C-peptide is also more immunogenically distinct between species with more available antibodies and a smaller target to increase the dynamic range of the anisotropy response. Finally, we used an N-terminally tagged C-peptide*. Our data showed that fluorophore in this position partially interferes with binding since the K_D1_ (180 ± 12 nM) was much larger than K_D2_ (5.8 ± 2 nM). Larger K_D_ values are often the result of faster off rates [22], which would effectively speed up the competitive binding kinetics [23]. Consistently, a C-terminal tagged C-peptide* with fluorophore further away from the epitope showed a much smaller K_D1_ and much slower equilibration time (**Fig. S1**). Overall, our design choices allowed us to detect the FAIA much closer to the trapped islet than we initially anticipated (i.e., in the 1^st^ pearl only ∼ 470 μm away from the islet) leading to lower dispersion and higher temporal resolution.

We used the InS-chip to measure insulin secretion from mouse islets in response to varying glucose and activators and inhibitors of the K_ATP_ channel. These data revealed a glucose-dependent double-peak of insulin secretion within the 1^st^ phase of both human and rodent islets [43]. We showed the time to reach the top of a single-peak response was similar to the time to reach the 2^nd^ peak of a double-peak response and that the double-peak response was more prevalent at high glucose stimulation. Thus, it seems the first peak results from higher metabolic flux. This may reflect glucokinase activity as the primary glucose sensor in β-cells, since previous work has shown small-molecule activators of glucokinase promoted earlier GSIS [44], [45]. To further demonstrate the InS-chip and explore the metabolism underlying 1^st^ phase insulin secretion, we imaged GSIS in parallel with MMP (i.e., TMRE) and cytosolic ATP levels (i.e., PercevalHR). Our results are consistent with glycolytic flux triggering secretion based on the high ATP response at low mitochondrial membrane potential (i.e., half-maximum MMP) [40]. Our results further suggest a transition to OxPhos generation of ATP in the subsequent peak since MMP reached its maximum response at this point in the response. Cationic TMRE sequesters into hyperpolarized mitochondria in response to active mitochondrial metabolism. The corresponding 2^nd^ peak of insulin secretion was consistently diminished in TMRE-loaded islets.

This artefact further implicates mitochondrial metabolism during the 2^nd^ peak of 1^st^-phase GSIS. Simultaneously, ATP recovered from a slight dip, which could reflect energy utilization due to exocytosis. In fact, Georgiadou et. al. showed that mitochondrial Ca^2+^ influx contributed to this ATP recovery, likely through activating various Ca^2+^-dependent dehydrogenases [38]. We envision this device could help further tease apart these mechanisms by coupling insulin secretion with live cell imaging of many other metabolic sensors.

The specific molecular events leading to a double-peaked secretion inside 1^st^-phase GSIS remain to be further elucidated, but a few models seem to be promising candidates. The first model depends on the metabolic and Ca^2+^ activity of individual β-cells within an intact islet. A small portion of β-cells (i.e., first responder or leader cells) lead the whole-tissue in cytoplasmic Ca^2+^ responses [46], [47]. Notably, the reported time delay between the fast and slow responders is similar to the time differences in the 1^st^ and 2^nd^ peaks within a double-peak response (i.e., a few mins). However, Ca^2+^ activity across the islet generally spreads more continuously thus it is unclear whether this model alone would result in two distinct peaks. The second model involves the secretion of different pools of insulin granules. It is at first tempting to incorporate the release of a readily releasable/docked pool of granules followed by the release of a reserve pool [48].

However, it is generally accepted that first-phase secretion involves docked granules and that the second phase involves the release of the reserve pool. More recently, the docked pool of insulin granules has been proposed to consist of two distinct populations: a slow-release population with low Ca^2+^ affinity and a fast-release population with high Ca^2+^ affinity. Yet the delay between the two pools seems too small to account for the difference in time between the first and second peaks. Finally, previous data suggest that b-cells are metabolically heterogeneous. Our data suggests the emergence of the first peak is triggered by glycolytic flux and the second peak is due to a transition to OxPhos, thus our data might reflect heterogeneity in this transition. In the end, these models are not mutually exclusive since the fusion of insulin granules with the plasma membrane is Ca^2+^-dependent [49].

The InS-chip has several limitations. First, our device was designed to hold a relatively narrow range of islet sizes (80 – 150 μm in diameter). Thus, we were unable to survey the full range of islet sizes to determine the effect on insulin secretion. A potential solution could be to incorporate islet traps with varying sizes and restrict upstream channel size accordingly to direct islets with different volumes. Second, our device did not accurately measure absolute insulin concentration. In part, we did this to maintain the simplicity of using a single pump to control flow. For example, we did not control nor account for varying flow rates in each of the separate chambers. Future work could explore particle image velocimetry to correct for varied flow rates in each channel.

Finally, our device has a relatively low throughput (4 islets/run) to ensure sufficient resolution of insulin secretion. The design could easily be scaled to accommodate more islets, but this would lead to progressively lower temporal resolution, particularly with simultaneous live cell imaging.

## Materials and Methods

### FAIA

Fluorescein-tagged C-peptide (C-peptide*) was purchased from Biomatik (Ontario, Canada). Stocks of the C-peptide* in 2% DMSO were stored at -80 °C and diluted in BMHH imaging buffer (125 mM NaCl, 5.7 mM KCl, 0.42 mM CaCl_2_, 0.38 mM MgCl_2_, 10 mM HEPES, and 0.1% BSA; pH 7.4) on the day of the experiment. The concentration of the tracer was quantified by absorbance of the FITC (ε = 73000 cm^-1^M^-1^) using a Nanodrop 2000 (ThermoFisher, United States). For the direct binding experiments, rat anti-mouse C-peptide monoclonal antibody (CII-29, NovusBio) was serially diluted in BMHH buffer with a fixed concentration of C-peptide* (100 nM). For competitive binding experiments, C-peptide (Biomatik, Canada) was serially diluted in BMHH buffer with a fixed concentration of C-peptide* (100 nM) pre-mixed with C-peptide antibody (200 nM). All assays were performed in Black 384-well plates with glass bottoms (CLS4581, Sigma Aldrich) with 10 μL per well at 37 °C using a custom-built wide-field fluorescence anisotropy microscope.

### Microfluidic device fabrication

Microfluidic devices were fabricated using soft lithography as described previously [28], [29]. Computer-Aided Design (CAD) files of the microfluidic devices were generated in AutoCAD (Autodesk, USA) for both fabrication and numerical simulation. A 120 μm thick layer of SU-8 2100 (Micro-Chem, USA) negative photoresist was spin-coated onto 6′′ silicon wafers (University Wafer) and exposed through a master printed on transparent plastic film (CAD/Art Services, USA). Devices were fabricated by pouring PDMS (Sylgard 184, Dow-Corning) at an elastomer base to curing agent mix poured over a master mold followed by degassing using a house vacuum and heating for 1 hr at 90 °C. The cured PDMS was peeled off, cut into pieces with individual device features, and hole-punched at inlets and outlets with a bevelled barrel of a 22G needle. The PDMS blocks were plasma bonded to no. 1.5 glass coverslips (VWR Scientific, USA). Circular PDMS wells (5 mm diameter; 4 mm tall) were plasma bonded on top of the inlets to act as media reservoirs as indicated.

### Microfluidic fluid flow modelling

Fluid flow in islet-on-a-chip was modelled with COMSOL Multiphysics 5.6 (COMSOL, Burlington, MA, USA) using the Laminar Flow module. Islets were modelled as solid 120 μm in diameter ellipsoids (i.e., porosity was omitted). For modelling purposes, pressure at the inlets of all devices was set to 0 Pa and fluid velocity at the outlets was set to values described in the text. The fluid property was based on water at 37 °C, which approximates the condition for empirical testing.

Shear stress on the islets was determined as the product of the shear rate and dynamic viscosity of water. A combination of *Laminar Flow* and *Transport of Diluted Species* was used to simulate mixing efficiency. C-peptide concentration in the outer channels was set to 200 nM and the diffusion coefficient was estimated to be 2.83 × 10^−11^ m^2^/s to set the lower range of the mixing efficiency [50]. The simulated models had a mesh with 874 391 to 919 856 triangular elements and 16045 to 30956 edge elements.

### Evaluation of mixing efficiencies of microfluidic devices

A microfluidic device was designed for the evaluation of mixing efficiencies of the reaction chamber. The device contains a three-channel inlet to simulate islet insulin secretion being sandwiched by carrier media. The device was flushed first with ethanol followed by BMHH buffer. 200 nM FICT-Dextran (FD150S, Sigma Aldrich) was placed in the outer two inlets while BMHH buffer was connected to the middle inlet. A pressure-based flow controller (Fluigent, US) was used to pull the solution from the single outlet at the indicated flow rate. Images of the microchannel were taken on a custom-built wide-field fluorescence anisotropy microscope.

Fluorescence intensity values across the microchannel were obtained in ImageJ using a line drawn perpendicularly to the direction of flow. Mixing efficiency at the region was determined based on [51]:

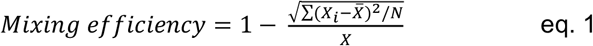

where, X_i_ is the fluorescence intensity of individual pixels across the channel, 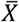 is the average of all fluorescence intensity and N is the total number of pixels.

### Particle streak velocimetry (PSV)

Images for PSV were collected using 40×/0.75 NA air objective (Olympus, Richmond Hill, Canada) on the custom-built wide-field fluorescence anisotropy microscope described below. Fluorescent beads with 1 μm diameter (FluoSpheres™ Carboxylate-Modified Microspheres) were diluted 500-fold in BMHH and drawn at 50 μL·h^-1^ with islets in the device. 10-20 sequential images were collected at the bypass channel and the channel leading to individual islet chambers as indicated using 505 nm excitation and 200 ms exposure. The apparent streak length caused by the movement of the beads during the exposure was measured in ImageJ and the linear velocity was calculated by dividing the streak lengths by the exposure time.

### Pancreatic islet isolation and culture

Animal procedures were approved by the Animal Care Committee of the University Health Network, Toronto, Ontario, Canada in accordance with the policies and guidelines of the Canadian Council on Animal Care (Animal Use Protocol #1531). Pancreatic islets were isolated from 10-12 weeks old FVB/ NJ male mice as previously described [28], [29] and cultured overnight in a humidified incubator (37°C, 5% CO_2_) unless stated otherwise.

### Imaging insulin secretion simultaneously with Ca^2+^ activity, mitochondrial membrane potential, and ATP

For Ca^2+^ imaging, islets were pre-incubated in BMHH with 4 μM Cal-520 (Life Technologies) for 1 hr for 30 min (37 °C, 5% CO_2_). For mitochondrial membrane potential imaging, islets were pre-incubated in BMHH with 20 nM Tetramethylrhodamine, ethyl ester (TMRE, MilliporeSigma) for 30 min (37 °C, 5% CO_2_). For ATP imaging, islets were infected with Perceval high range (HR) adenovirus under the control of the rat insulin promoter (a gift from Matthew Merrins). Twenty freshly isolated mouse islets per run were infected with 10^7^ PFU/mL for 1 hr on a Belly Dancer

Shaker followed by an additional 18 hr in a humidified incubator (37°C, 5% CO_2_). Media was subsequently replaced with fresh culture media and islets were incubated for an additional 24 hr. The labelled islets were loaded into a stage-mounted islet-on-a-chip device (37 °C, 50 μL·h^-1^). Islets were exposed to different treatments as indicated by switching media in the on-chip reservoir. A custom macro was used to image C-peptide* (500 nm, 1.5-1.8 s exposure) at 10 s intervals simultaneously with Cal520 (505 nm, 0.2 s exposure), TMRE (535 nm, 0.3 s exposure), or PercevalHR (505 nm, 0.2 s exposure).

### Widefield Fluorescence Anisotropy Microscope

Standard fluorescence and steady-state fluorescence anisotropy imaging were done using an inverted widefield RAMM fluorescence microscope (ASI) [36], [52]. The microscope is equipped with 3 LEDs (405/465, 505/535, and 590 nm) and a stage top incubator (Okolab). A 40×/0.75 NA air objective (Olympus, Richmond Hill, Canada) was used for all experiments. For standard fluorescence, the emission was passed through an eYFP (ET535/30M) emission filter on a filter wheel and collected by an Iris 15 sCMOS camera (Photometrics). For studies coupling fluorescence and steady-state fluorescence anisotropy imaging, the excitation light was linearly polarized prior to exciting the sample. To image TMRE intensity and C-peptide* anisotropy, the sample was excited using the 535 nm and 465 nm LED, while emission was collected through 590-660 nm and 495-525 nm band-pass emission filters, respectively. To image PercevalHR intensity and C-peptide* anisotropy, the sample was excited using the 505 nm LED, while emission was collected through 495-525 nm band-pass emission filter, respectively. For both, the filtered emission light was split using an Optosplit II (Cairn, Faversham, UK) to simultaneously collect parallel (*I*_*ǁ*_) and perpendicular (*I*_⊥_) light on separate regions of the camera.

## Statistical analysis

Parallel (*Iǁ*) and perpendicular (*I*⊥) fluorescence intensity images were analyzed with a custom ImageJ plugin that calculated the pixel-by-pixel anisotropy (r) based on [53]:

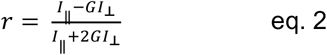

Here, G is a factor used to account for the skewing of the anisotropy by the microscope. Blank buffer was imaged at the same intensity and exposure to determine the background. We measured the G-factor using a solution of 100 nM fluorescein, which has a near-zero steady-state fluorescence anisotropy simplifying Eq. 2 to G = *I*_*ǁ*_ */ I*_⊥_ [54].

The measured fluorescence anisotropy can be related to the fraction of bound tracer (F_SB_) through [21]:

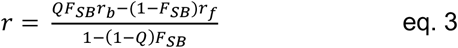

Here, Q is the ratio of fluorescence intensity of bound to free tracer, and r_b_ and r_f_ are the anisotropies of bound and free tracer, respectively. The average value of Q used in the fitting (0.822) was determined based on images of the assay mixture.

K_D1_ and K_D2_ were subsequently fitted in OriginLab (Northampton, MA) using equations described by Roehrl et al. [21]:

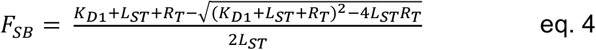

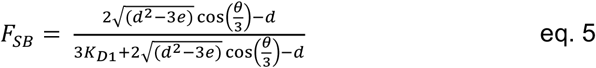

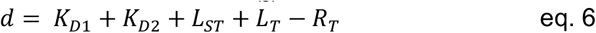

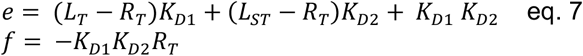

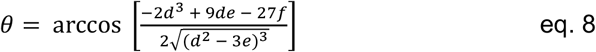

Here, L_ST_, L_T_ and R_T_ are the total concentration of the fluorescent probe, the full-length C-peptide competitor, and the antibody, respectively. Eq. 3-4 were used to fit for K_D1_, while eq. 3, 5-8 was used to fit for K_D2_.

## Acknowledgments

This work was supported by grants from NSERC (RGPIN-2022-04454 and DGDND-2022-04454) and CIHR (CIHR PJT-162330) to JVR. Stipend support was provided by Mary H. Beatty Fellowship to YW and NSERC Postgraduate Scholarship Doctoral (NSERC PGSD-535059-2019) to RR.

