## Supplemental Figures for "Insulin secretion chip (InS-chip) reveals two peaks within first-phase that temporally coincide with a transition in glucose metabolism"

**Supporting Information for**

### Design of a fluorescence anisotropy immunoassay on-a-chip to measure insulin secretion (InS-chip) from individual islets with coupled live cell imaging

Yufeng Wang, Romario Regeenes, Mahnoor Memon, and Jonathan V. Rocheleau

* Jonathan V. Rocheleau

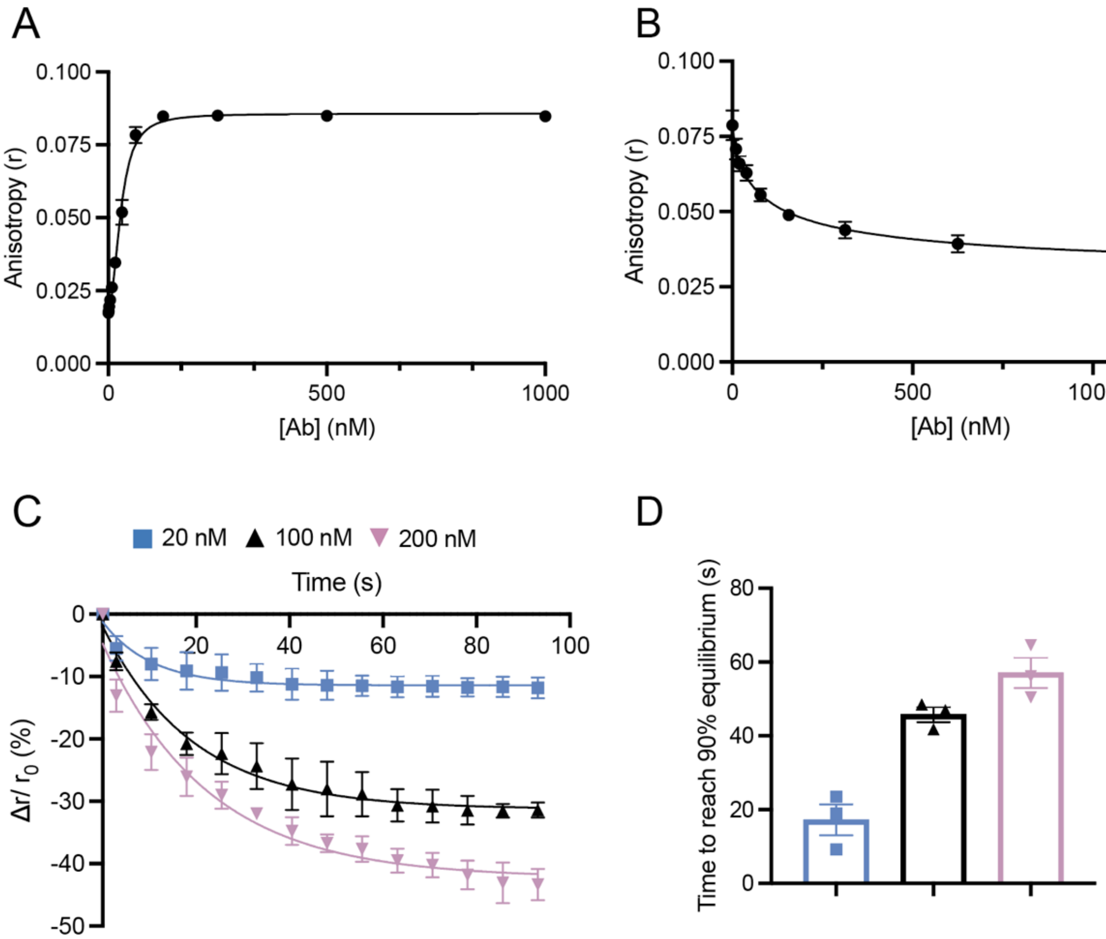


**Fig. S1. Characterization of FAIA using a high-affinity C-terminal-tagged version of C-peptide*.** (**A**) Fitting of the direct binding curves between serially diluted Ab and 100 nM C-peptide* yielded a $K_{D1}$ of 2.89 ± 1.44 nM (n = 3). (**B**) Fitting of the competitive binding curves between serially diluted C-peptide and pre-mixed 30 nM Ab and 100 nM C-peptide* yielded a $K_{D2}$ of 4.12 ± 1.6 nM (n = 3). (**C**) Kinetics of competitive binding between different concentrations of C-peptide and pre-mixed 30 nM Ab and 100 nM C-peptide* in the device shown in Fig. S3A (n = 3 for each C-peptide concentration). (**D**) The time to reach the 90% plateau of the competition kinetics curves is shown in (**C**).

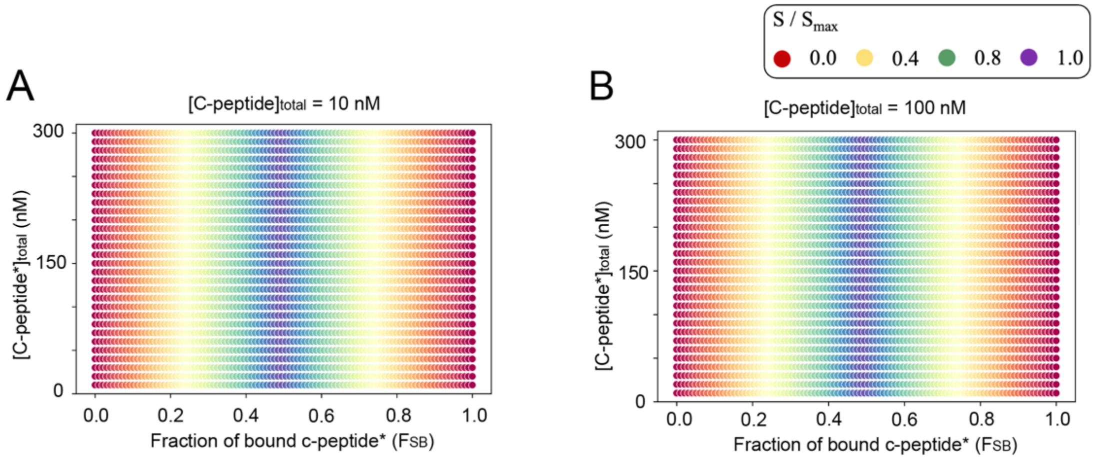


**Fig. S2. Sensitivity optimization reveals maximum sensitivity is attained at 50% bound.** Sensitivity was calculated by keeping antibody concentration, $K_{D1}$ and $K_{D2}$ constant while varying fraction bound, the total concentration of C-peptide* and unlabelled C-peptide. The sensitivity rank-plot reveals the 50% bound state is the most sensitive when competing with (**A**) 10 nM and (**B**) 100 nM of unlabelled C-peptide.

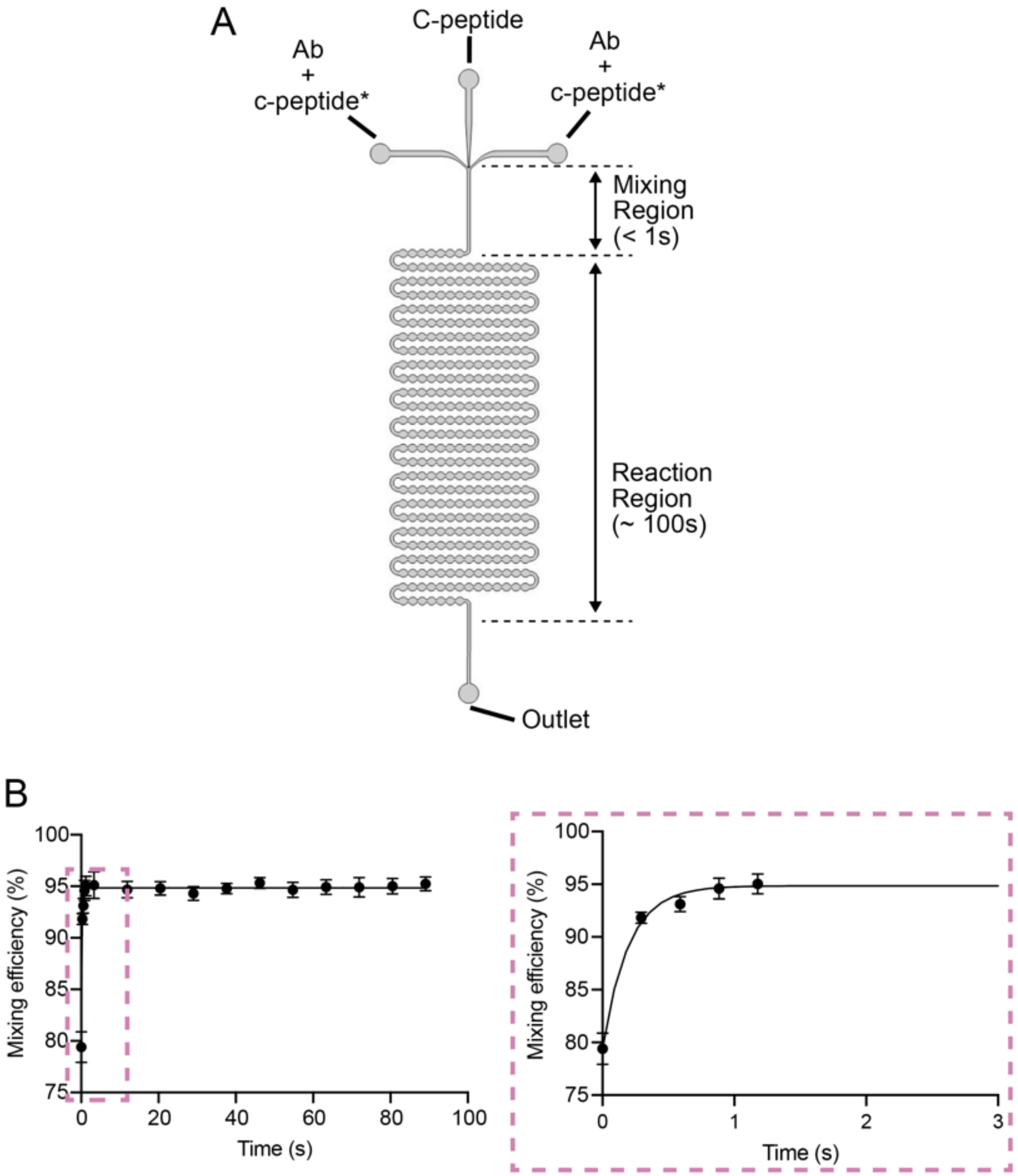


**Fig. S3. Microfluidic device designed to measure the temporal dynamics of the FAIA.** (**A**) Schematic of the device that simulates a continuous-flow method for studying assay kinetics. The device consists of a straight 100 μm wide channel for fast mixing of assay reagents (Mixing region) and a serpentine channel for monitoring the progress of competitive binding of the FAIA (Reaction region). (**B**) The average mixing efficiency along the microfluidic device was measured using 200 nM FITC-Dextran (~150 kDa) drawn from two outer inlets sandwiching clear imaging media in the middle inlet at a total flow rate 25 μL/hr (n = 5). Insert graph (dashed box) shows the change in mixing within the first 2 s.

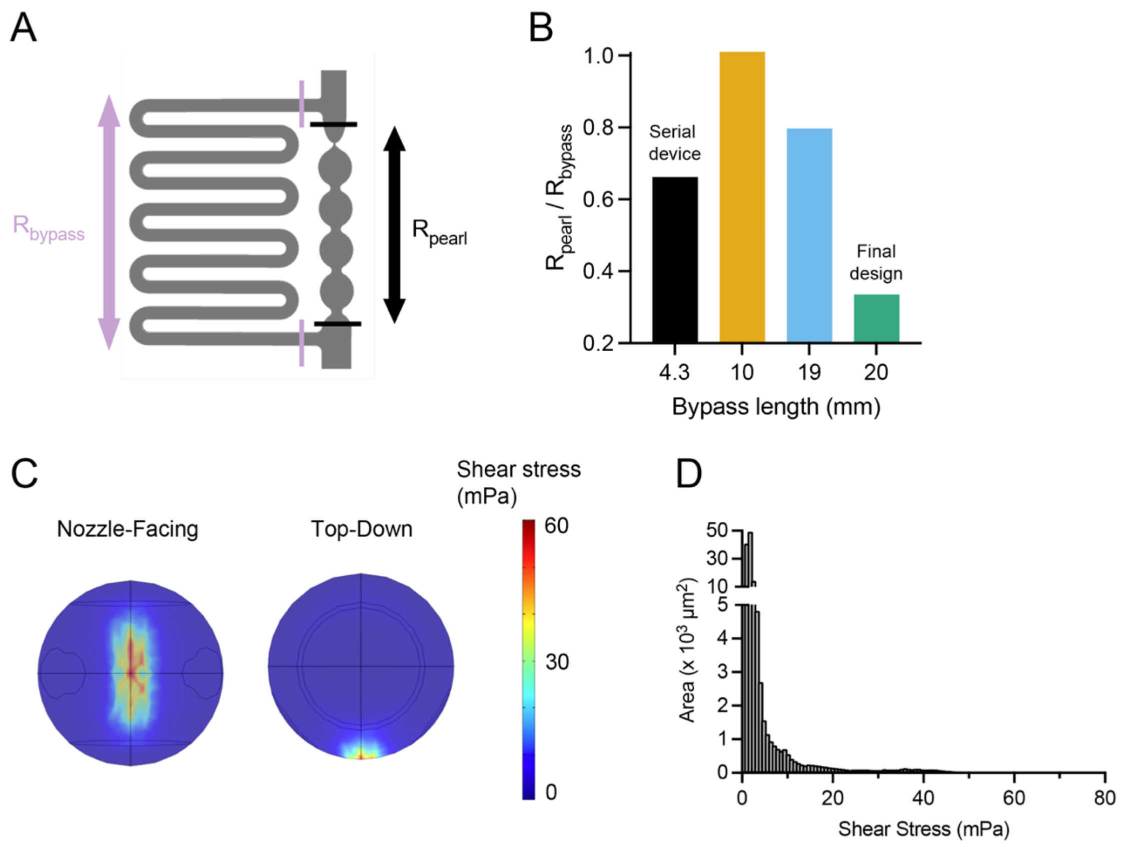


**Fig. S4. Characterization of fluid dynamics in the islet-on-a-chip.** (**A**) Schematic of a single islet chamber to illustrate different segments of channels used for calculation of hydrodynamic resistance. (**B**) The ratio of the resistance from the pearl channel to the bypass channel among different designs based on Laminar Flow simulation in COMSOL. (**C**) Shear stress induced on a loaded islet in the islet-on-a-chip based on Laminar Flow simulation. Images demonstrate nozzle-facing (*left*) and top-down (*right*) views of the simulated islet. (**D**) The distribution of shear stress induced on the islet reveals that most of the surface experiences less than 10 mPa of shear stress with a flow rate of 50 $\mu L \cdot h^{-1}$.

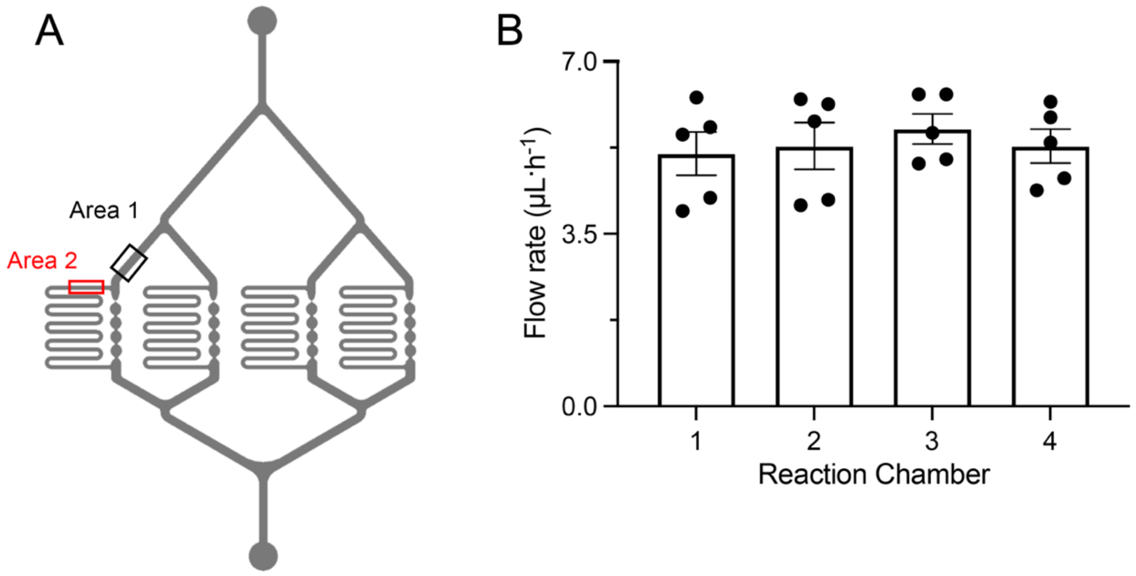


**Fig. S5. Measurement of flow rate in the reaction chamber with islets loaded in the islet-on-a-chip.** (**A**) Imaging areas for flow rate measurement in the straight channel leading to an individual islet chamber (Area 1) and the bypass channel of the same islet chamber (Area 2). (**B**) The device was loaded with islets and flow rates in Area 1 and 2 were measured with fluorescent beads using particle streak velocimetry. Beads were imaged with 505 nm excitation and 200 ms exposure. The flow rate in the reaction chamber downstream of the loaded islet trap was estimated by subtracting the flow rate measured in Area 2 from that in Area 1 (n = 5).

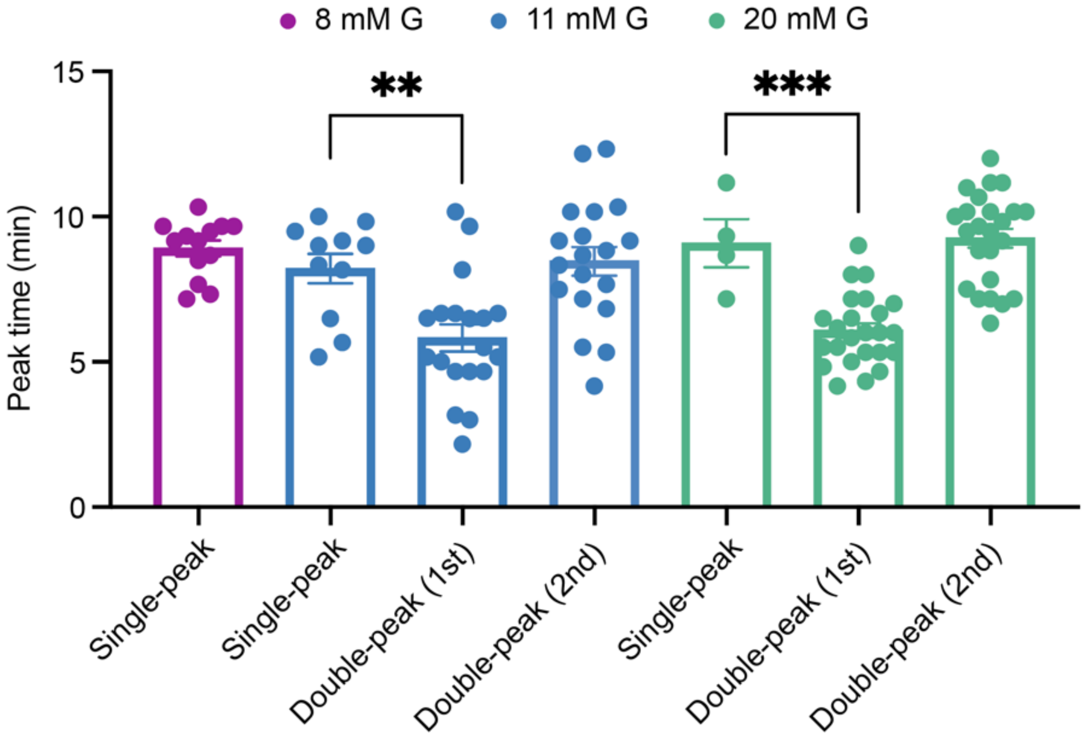


**Fig. S6. Characterization of peak time of 1st phase GSIS.** The peak time (i.e., local maximum) for the GSIS traces (representative traces shown in **Fig. 6A**) with single- and double-peak 1st phase response was compared. All GSIS traces for 8 mM glucose (n = 13) contained single peaks, while both 11 mM (n = 30) and 20 mM glucose (n = 28) showed either single- or double peaks. ** indicated $p < 0.01$ and *** indicates $p < 0.001$ by unpaired t-test.
